# Nucleotide Archival Format (NAF) enables efficient lossless reference-free compression of DNA sequences

**DOI:** 10.1101/501130

**Authors:** Kirill Kryukov, Mahoko Takahashi Ueda, So Nakagawa, Tadashi Imanishi

## Abstract

**Summary:** DNA sequence databases use compression such as gzip to reduce the required storage space and network transmission time. We describe Nucleotide Archival Format (NAF) – a new file format for lossless reference-free compression of FASTA and FASTQ-formatted nucleotide sequences. NAF compression ratio is comparable to the best DNA compressors, while providing dramatically faster decompression. We compared our format with DNA compressors: DELIMINATE and MFCompress, and with general purpose compressors: gzip, bzip2, xz, brotli, and zstd.

**Availability:** NAF compressor and decompressor, as well as format specification are available at https://github.com/KirillKryukov/naf. Format specification is in public domain. Compressor and decompressor are open source under the zlib/libpng license, free for nearly any use.

## Introduction

DNA sequence databases are growing exponentially, owing to the continuing advances in sequencing technologies. Data compression is typically used for all stored DNA sequence data to save storage space and network transmission times. In 1993 the first specialized DNA compressor was proposed (Grumbach and Tahi, 1993). Since then, numerous DNA compressors were developed (e.g., Cao et al., 2007, Li et al., 2013, Benoit et al., 2015, Al-Okaily et al., 2017). In our experience only two compressors pass the practicality threshold: DELIMINATE (Mohammed et al., 2012) and MFCompress (Pinho and Pratas, 2014). They are stable, support commonly used features of FASTA format, and are efficient enough to be able to handle practical tasks such as compressing (and decompressing) entire vertebrate genomes. Although they achieve impressive compression ratios, both DELIMINATE and MFCompress have very slow decompression, significantly limiting their usefulness with large databases.

Despite the numerous studies on DNA compression, currently the majority of sequence databases continue to rely on gzip (https://www.gzip.org/). We attribute this enduring popularity to gzip’s wide availability, robustness, and speed (especially decompression speed). These qualities appear to be able to outweigh gzip’s less than stellar compression ratio. Other popular general purpose compressors have been developed, such as bzip2 (http://www.bzip.org/) and lzma/xz (https://tukaani.org/xz/format.html). In addition, recent years have seen the emergence of a new generation of advanced general purpose compressors, most notably brotli (https://github.com/google/brotli) and zstd (https://github.com/facebook/zstd). These compressors improve upon gzip performance, but still cannot touch specialized DNA compressors in compression strength.

In this work we describe a new DNA compression format called Nucleotide Archival Format (NAF). NAF provides a state of the art compression ratio, on par with DELIMINATE and slightly behind MFCompress. At the same time, it provides 30 to 80 times faster decompression. NAF compresses and decompresses from/to FASTA and FASTQ for-mats. NAF supports masked sequence and ambiguous IUPAC nucleotide codes. NAF has no restrictions on sequence length, and does not require reference sequences.

## Methods

Analogously to many previous methods, including DELIMINATE and MFCompress, NAF compression operates by splitting the input into headers, nucleotide sequences, mask (in case of masked sequence), and qualities (in case of FASTQ input), which are processed separately. Sequences are concatenated together, with lengths stored separately. The combined nucleotide sequence is then converted into 4-bit encoding, which stores 2 nucleotides per byte. This representation is extremely fast, for both encoding and decoding, and allows natively representing ambiguous IUPAC nucleotide codes (NC-IUB, 1985), including ‘N’, ‘Y’, ‘R’, etc. All the resulting data streams are then compressed with the general purpose compressor zstd.

The decompression consists of decompressing those separate streams, and re-assembling them together into FASTA or FASTQ-formatted output. NAF’s high decompression speed owes to these factors: 1) Using zstd compressor, which itself is designed for fast decompression. 2) In NAF, zstd works with 4-bit encoded sequence, which means that it deals with data half the size of original sequence. 3) Decoding of 4-bit sequence is very fast using a simple table lookup for pairs of nucleotides.

NAF implementation provides interface that is friendly to automated sequence analysis workflows. NAF compressor reads data from standard input stream, enabling on-the-fly compression of data originating from other process. Similarly, NAF decompressor allows piping decompressed sequences directly into the next analysis step. In addition, NAF allows decompressing only headers or only sequences, as well as 4-bit encoded sequence. This will allow applications such as sequence search or composition analysis to work directly with 4-bit encoded sequence.

## Results

We compared NAF with DNA compressors DELIMINATE and MFCompress, as well as general purpose compressors: gzip, bzip2, xz, brotli, and zstd. Figs. 1, S1, S2 and Table S1 show their results on the human genome. Fig. S3 and Table S2 show overall compression ratios on larger set of genomes. Table S3 shows the result on FASTQ data. Table S4 compares availability and features of these compressors.

**Fig. 1.**
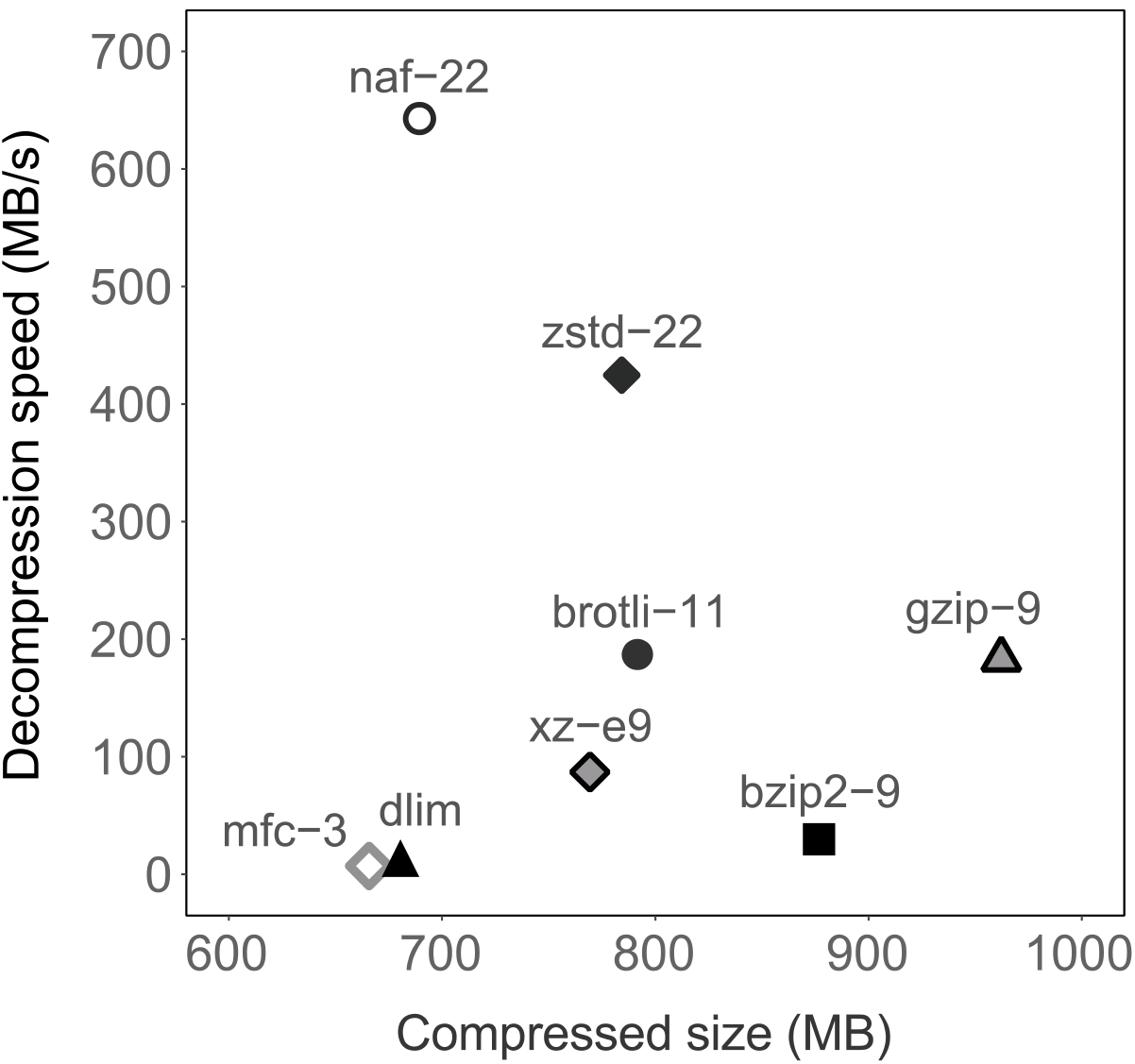
Compression strength and decompression speed of 8 compressors. Human genome (GRCh38, 3.3 GB) was used as test dataset. “mfc” and “dlim” represent MFCompress and DELIMINATE, respectively. Each compressor was used with its strongest compression setting: ‘gzip -9’, ‘bzip2 -9’, ‘brotli -11’, ‘zstd --ultra -22’, ‘xz -e9’, ‘ennaf -22’, ‘delim a’, ‘MFCompressC -3’. CPU used: Intel Xeon E5-2643v3 (3.4 GHz).

Compression strength of NAF is close to DELIMINATE and slightly behind MFCompress. All three DNA compressors achieve markedly better compression ratio than the general purpose compressors. However what sets NAF apart is its high speed of decompression. In the human genome example, NAF decompression is faster by a factor of 35 and 78 than DELIMINATE and MFCompress, respectively. Also NAF provides high compression speed in its fast mode, using “−1” option of the compressor (Fig. S2, Tables S1 and S3).

When considering the typical application of compression for distributing data from sequence databases, we can estimate the total time that it takes from initiating download until accessing the decompressed data, for different compressors. Fig. S1 compares access times for 8 compressors, as well as for the uncompressed FASTA format, while assuming link speeds of 100 Mbit/s and 1000 Mbit/s. It can be seen that NAF enables the fastest distribution of data over network, allowing to reduce waiting time and bandwidth cost (in addition to reducing the required storage space).

In Fig. S2 we compared the total transfer time including compression, for a hypothetical scenario of one-time data transfer. In this case the fast setting (“−1” option) of NAF provides the best total transfer time. As Tables S1 and S3 show, NAF’s fast mode retains useful compression strength, which is still better than gzip’s strong mode (“−9”).

## Conclusion

NAF offers significant advantages over both general purpose and specialized DNA compressors. NAF’s combination of compactness and decompression speed enables rapid distribution of database sequences to users. NAF’s fast decompression allows storing NAF-compressed databases, decompressing them on-the-fly when necessary. NAF also provides a useful combination of compactness and high compression speed in fast mode, making it ideal for one-time data transfer. We believe that our new format will save network bandwidth, time, and disk space, and thus contribute greatly to both operators and users of DNA sequence databases.

