## Supplementary Figures and Tables for "Nucleotide Archival Format (NAF) enables efficient lossless reference-free compression of DNA sequences"

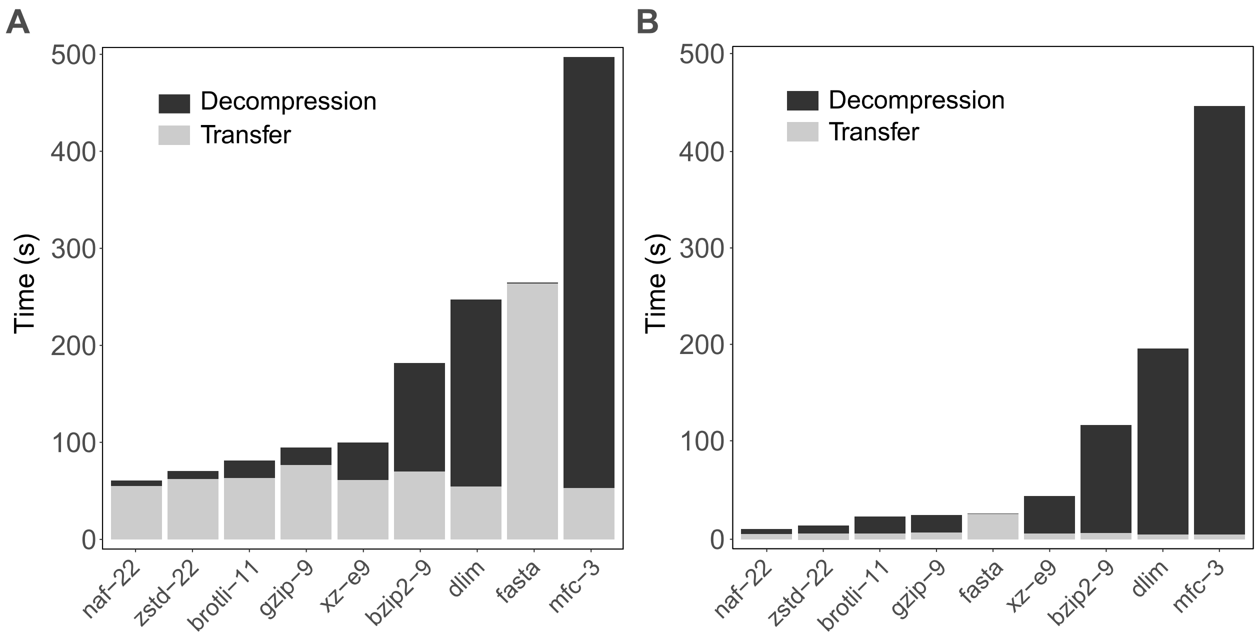


**Fig S1.** Time of accessing the human genome (GRCh38, 3.3 GB) stored on a remote server, in various formats (including the uncompressed FASTA format). Total time consists of network transfer time (estimated for a link speed of 1000 Mbit/s) and decompression time (measured). Panel A assumes network speed of 100 Mbit/s, B - 1000 Mbit/s. This figure shows the strongest compression setting of each compressor, as normally used in sequence databases.

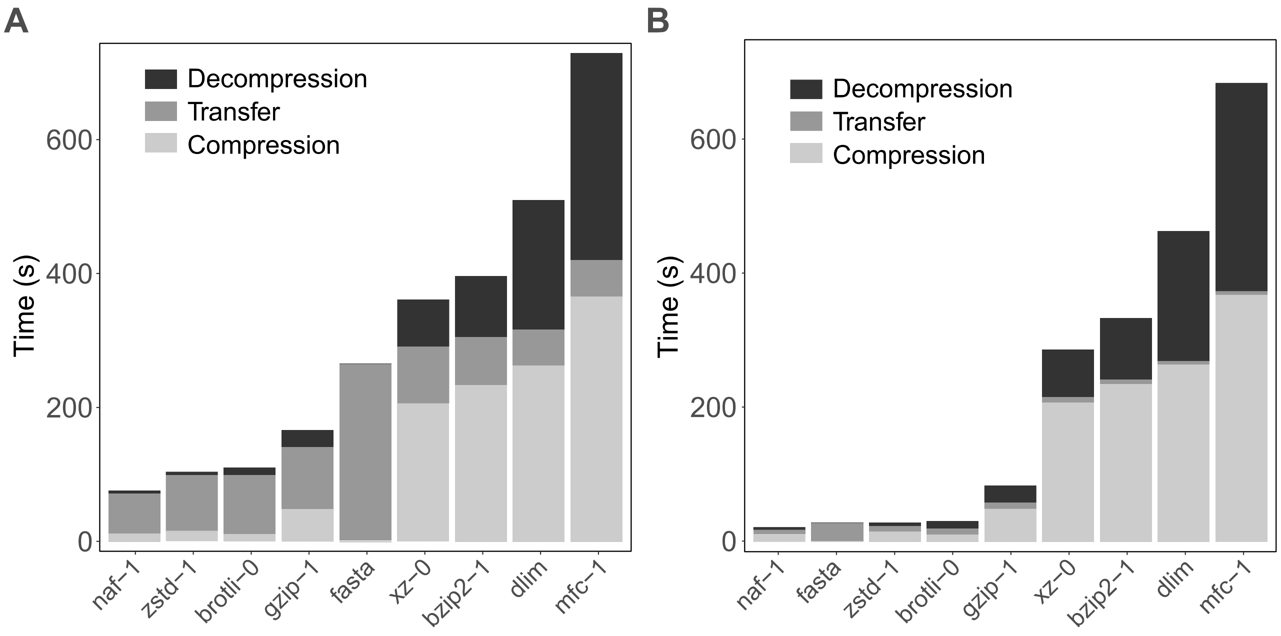


**Fig S2.** Time of transferring the human genome (GRCh38, 3.3 GB) over network, in various formats (including the uncompressed FASTA format). Total time consists of compression time (measured), network transfer time (estimated) and decompression time (measured). Panel A assumes network speed of 100 Mbit/s, B - 1000 Mbit/s. This figure uses the fastest setting of each compressor, typically used for one-time data transfer.

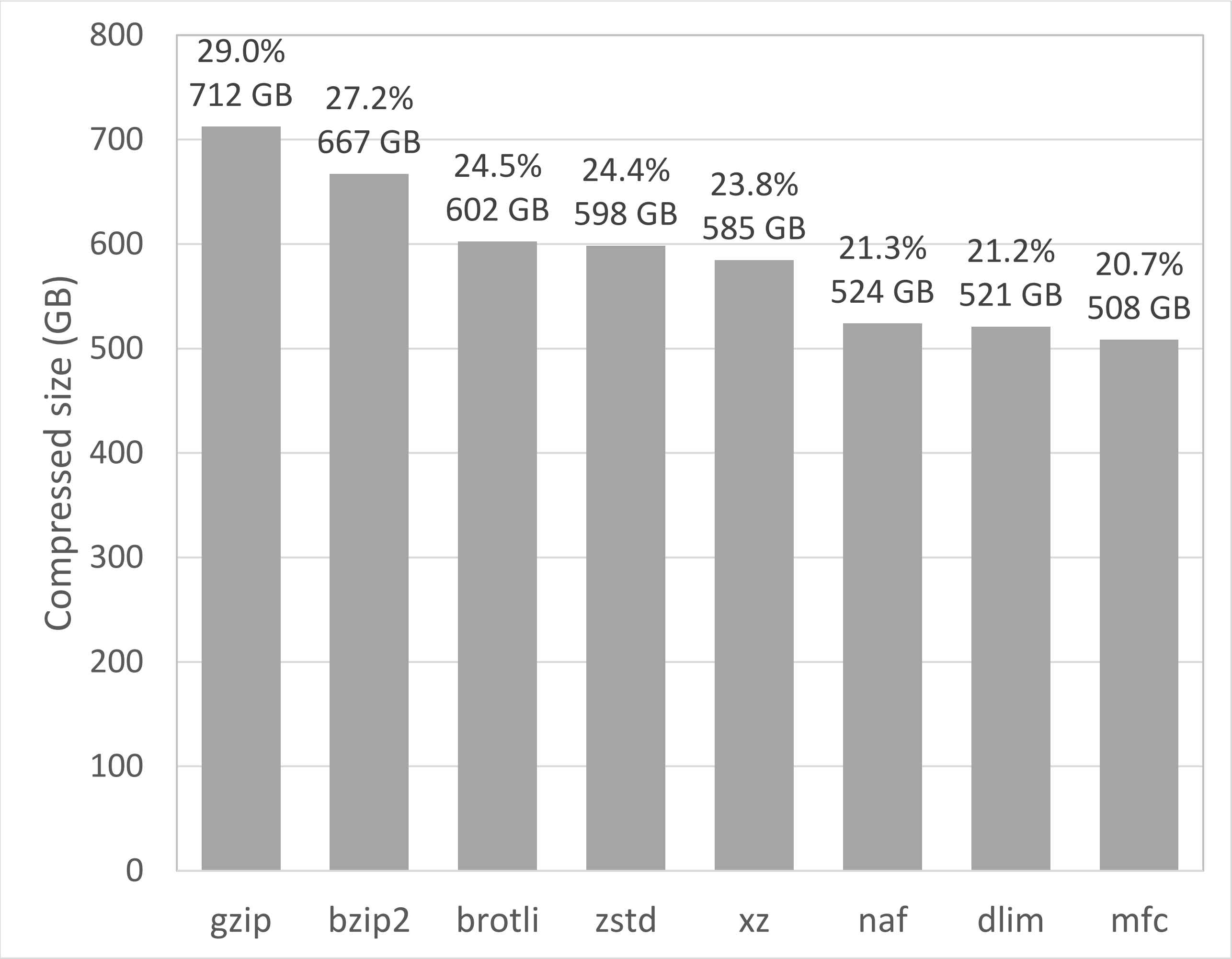


**Fig. S3.** Comparison of genome compression strength between 8 compressors. A collection of genome sequences obtained from GenomeSync (http://genomesync.org/) was used as test data. GenomeSync snapshot from 2018-11-23 was used, containing 196,199 genomes, with total FASTA file size of 2,457 GB. Each genome was compressed separately, so 196,199 files were produced with each compressor. "dlim" means DELIMINATE, "mfc" means MFCompress. Percent ratio is computed as (compressed size / FASTA size x 100). Compressor options used: 'gzip -9', 'bzip2 -9', 'brotli -11', 'zstd --ultra -22', 'xz -e9', 'ennaf -22', 'delim a', 'MFCompressC -3'.

**Table S1.** Compressed size, compression and decompression time on the human genome (GRCh38, 3.3 GB). CPU used: Intel Xeon E5-2643v3 (3.4 GHz). The 2-nd decompression run was timed, so that all compressed files were in disk cache, and disk speed did not influence the result. This table includes both the fastest and the strongest setting of each compressor.

| Compressor | Compressed size (bytes) | Compression time (s) | Decompression time (s) |
|---|---|---|---|
| gzip-1 | 1,156,039,726 | 49.0 | 25.4 |
| gzip-9 | 960,058,842 | 1,037.7 | 17.9 |
| bzip2-1 | 902,487,201 | 233.8 | 90.5 |
| bzip2-9 | 875,294,078 | 256.4 | 111.6 |
| xz-0 | 1,058,102,156 | 206.4 | 70.4 |
| xz-e9 | 766,919,800 | 4,150.7 | 38.6 |
| brotli-0 | 1,100,086,261 | 11.2 | 10.5 |
| brotli-11 | 792,713,109 | 9,043.1 | 17.5 |
| zstd-1 | 1,040,619,629 | 15.7 | 4.9 |
| zstd-22 | 784,452,563 | 4,859.9 | 7.8 |
| **naf-1** | **753,880,552** | **12.0** | **3.9** |
| **naf-22** | **688,685,727** | **1,950.4** | **5.1** |
| mfc-1 | 687,853,201 | 366.0 | 308.5 |
| mfc-3 | 664,726,085 | 634.4 | 443.9 |
| dlim | 680,899,418 | 262.8 | 192.6 |

**Table S2.** Compressed size and decompression time on a set of genomes of fingi, algae and protists. Genomes were obtained from GenomeSync (http://genomesync.org/): snapshot from 2018-11-23, 4,662 genomes, 168 GB in FASTA format. 1 genome per file was stored in each format. CPU used: Intel Xeon E5-2643v3 (3.4 GHz). Storage: RAID 0 array of 4 SSDs. All data was not in disk cache prior to decompression, so decompression time includes time of reading the files. For FASTA, the time of "cat >/dev/null" was recorded. Strongest compression setting was used for each compressor.

| Compressor | Compressed size (bytes) | Decompression time (s) |
|---|---|---|
| fasta | 168,143,459,104 | 210.1 |
| gzip-9 | 50,836,404,143 | 1,076.8 |
| bzip2-9 | 47,322,681,943 | 5,902.9 |
| brotli-11 | 43,570,617,258 | 1,250.3 |
| zstd-22 | 43,140,888,642 | 457.7 |
| xz-e9 | 42,266,241,432 | 1,990.4 |
| dlim | 38,517,371,519 | 12,103.0 |
| **naf-22** | **38,398,633,866** | **331.6** |
| mfc-3 | 37,141,175,505 | 58,936.7 |

**Table S3.** Compressed size and decompression time on FASTQ dataset DRR000020 (https://www.ncbi.nlm.nih.gov/sra/?term=DRR000020), 264 MB in FASTQ format. CPU used: Intel Xeon E5-2643v3 (3.4 GHz). The 2-nd decompression run was timed, so that all compressed files were in disk cache, and disk speed did not influence the result. This table includes both the fastest and the strongest setting of each compressor.

| Compressor | Compressed size (bytes) | Compression time (s) | Decompression time (s) |
|---|---|---|---|
| gzip-1 | 120,406,516 | 4.3 | 2.3 |
| gzip-9 | 105,814,822 | 40.4 | 1.7 |
| bzip2-1 | 93,680,349 | 18.9 | 8.3 |
| bzip2-9 | 88,554,581 | 20.4 | 9.8 |
| xz-0 | 112,267,112 | 19.9 | 6.7 |
| xz-e9 | 64,679,088 | 311.4 | 3.1 |
| brotli-0 | 119,430,476 | 1.2 | 1.1 |
| brotli-11 | 71,296,854 | 655.7 | 0.9 |
| zstd-1 | 115,533,089 | 1.3 | 0.4 |
| zstd-22 | 64,477,941 | 313.8 | 0.6 |
| **naf-1** | **91,086,071** | **1.3** | **0.6** |
| **naf-22** | **59,966,838** | **207.1** | **0.8** |

**Table S4.** Availability and features of the 8 compressors.

| Compressor | gzip | bzip2 | xz | brotli | zstd | dlim | mfc | naf |
|---|---|---|---|---|---|---|---|---|
| Free for non-commercial use | ✓ | ✓ | ✓ | ✓ | ✓ | ✓ | ✓ | ✓ |
| Free for commercial use | ✓ | ✓ | ✓ | ✓ | ✓ | ✗ | ✗ | ✓ |
| Source available | ✓ | ✓ | ✓ | ✓ | ✓ | ✗ | ✓ | ✓ |
| Format specification available | ✓ | ✓ | ✓[1] | ✓ | ✓ | ✗ | ✗ | ✓ |
| Web-site available | ✓ | ✓ | ✓ | ✓ | ✓ | ✗[2] | ✓ | ✓ |
| Supports FASTA | ✓ | ✓ | ✓ | ✓ | ✓ | ✓ | ✓ | ✓ |
| Supports FASTQ | ✓ | ✓ | ✓ | ✓ | ✓ | ✗ | ✗ | ✓ |
| Pipe-in for compression | ✓ | ✓ | ✓ | ✓ | ✓ | ✗ | ✗ | ✓ |
| Pipe-out for decompression | ✓ | ✓ | ✓ | ✓ | ✓ | ✗ | ✗ | ✓ |
| Can specify output file name for compression | ✓ | ✓ | ✓ | ✓ | ✓ | ✗[3] | ✓ | ✓ |

[1] Unofficial specification.

[2] Binary is available by emailing the authors.

[3] Automatically names output file by taking input file name and appending ".dlim" to it.
